# Mitochondrial sirtuin TcSir2rp3 affects TcSODA activity and oxidative stress response in *Trypanosoma cruzi*

**DOI:** 10.1101/2021.09.06.458705

**Authors:** Leila dos Santos Moura, Vinícius Santana Nunes, Antoniel A. S. Gomes, Ana Caroline de Castro Nascimento Sousa, Marcos R. M. Fontes, Sergio Schenkman, Nilmar S. Moretti

**Author notes:** Correspondence to Nilmar S. Moretti.

## Abstract

*Trypanosoma cruzi*, the etiological agent of Chagas disease, faces a variety of environmental scenarios during its life cycle in both invertebrate and vertebrate hosts, which include changes in the redox environment that requires a fine regulation of a complex antioxidant arsenal of enzymes. Reversible post-translational modifications, as lysine acetylation, are a fast and economical way for cells to react to environmental conditions. Acetylation neutralizes the lysine positive charge conferring novel properties to the modified proteins, from changes in enzymatic activity to subcellular localization. Recently, we found that the main antioxidant enzymes, including the mitochondrial superoxide dismutase A (TcSODA) are acetylated *in T. cruzi*, suggesting that protein acetylation could participate in the oxidative stress response in *T. cruzi*. Therefore, we investigated whether mitochondrial lysine deacetylase sirtuin 3 (TcSir2rp3) was involved in the activity control of TcSODA. We observed an increased resistance to hydrogen peroxide and menadione two oxidant compounds in parasites overexpressing TcSir2rp3. Increased resistance was also found for benznidazole and nifurtimox, the two drugs available for treatment of Chagas disease, known to induce reactive oxidative and nitrosactive species in the parasite. In parallel, TcSir2rp3 overexpressing parasites showed parasites showed a reduction in the ROS levels after treatment with benznidazole and nifurtimox, suggesting a role of TcSir2rp3 in the oxidative stress response. To better understand the way TcSir2rp3 could contributes to oxidative stress response, we analyzed the expression of a key antioxidant enzyme, TcSODA, in the TcSir2rp3 overexpressing parasites and did not detect any increase in protein levels of this enzyme. However, we found that parasites overexpressing TcSir2rp3 presented higher levels of superoxide dismutase activity, and also that TcSir2rp3 and TcSODA interacts in vivo. Knowing that TcSODA is acetylated at lysine residues K44 and K97, and that K97 is located at similar region in the protein structure as K68 in human manganese superoxide dismutase (MnSOD), responsible to regulates MnSOD activity, we generated mutated versions of TcSODA at K44 and K97 and found that replacing K97 by glutamine, which mimics an acetylated lysine, negatively affects the enzyme activity in vitro. By using molecular dynamics approaches we revealed that acetylation of K97 induces specific conformational changes in TcSODA with respect of hydrogen bonding pattern to neighbor residues, specifically D94 and E96, suggesting a key participation of this residue to modulate the affinity to O_2_- by changing the charge availability on the surface of the enzyme. Taken together, our results showed for the first time the involvement of lysine acetylation in the maintenance of homeostatic redox state in trypanosomatids, contributing to the understanding of mechanisms used by *T. cruzi* to progress during the infection and opening the opportunity to explore protein acetylation as potential drug target in this parasite.

## Introduction

*Trypanosoma cruzi* is a protozoan parasite of the Trypanosomatidae family responsible for Chagas disease, an illness endemic in 21 countries across Latin America and now being detected in the southern USA and many other countries. Around 7 million people are infected and more than 70 million live at risk to be infected in the world (Perez-Molina and Molina, 2018;Lidani et al., 2019;Moretti et al., 2020). The parasite has a heteroxenous life cycle shifting between an invertebrate and a vertebrate host. Metacyclic trypomastigote parasites are transmitted by triatomine bugs feces during blood meal and can infect any nucleated cell type. Inside the vertebrate host cell, the metacyclic trypomastigotes differentiate to amastigote replicative forms, which latter on will differentiates to trypomastigotes that will be released upon cell lyses and can infect other cells or be ingested by triatomine bugs during a bloodmeal. In the invertebrate host, trypomastigotes differentiate to replicative epimastigotes that in the hindgut will generate metacyclic trypomastigotes, which can infect another vertebrate host (Perez-Molina and Molina, 2018;Moretti et al., 2020). During its life cycle transitions *T. cruzi* faces variable environmental conditions, such as, changes in temperature, nutritional state, oxidant products, and the parasite must adapt to these alterations to survive and succeed in the infection (Moretti and Schenkman, 2013).

Reversible post-translational modifications (PTMs) are a fast and economical way for cells to react to environmental conditions. PTMs such as methylation, phosphorylation and acetylation, are detected on hundreds of proteins in the cells (Choudhary et al., 2014;Levy, 2019;Macek et al., 2019). Acetylation of lysine residues is one of the most common PTMs and it is characterized by the addition of an acetyl group to the ε-amino group of this protein residue. The acetyl group neutralizes the lysine positive charge conferring novel properties to the modified proteins, from changes in enzymatic activity to subcellular localization (Narita et al., 2019).

This PTM has been identified in several prokaryote and eukaryote species (Carabetta and Cristea, 2017;Narita et al., 2019;Maran et al., 2021). Recently, we have described the acetylome, set of lysine-acetylated proteins, of procyclic and bloodstream forms of *T. brucei*, and of *T. cruzi* epimastigotes (Moretti et al., 2018). We detected 288 lysine acetylation sites (Kac) in 210 proteins of procyclic form, and 380 Kac in 285 proteins in the bloodstream form, and in *T. cruzi* epimastigotes we found 389 Kac sites in 235 proteins (Moretti et al., 2018). Interesting, in *T. brucei* glycolytic enzymes were found heavily acetylated and later proved to be regulated by acetylation (Barbosa Leite et al., 2020), while in *T. cruzi* several proteins that are central for antioxidant defense of the parasite were detected acetylated (Moretti et al., 2018).

The addition and removal of acetyl groups on lysines are coordinated by lysine acetyltransferases (KATs) and lysine deacetylases (KDACs), respectively (Marmorstein and Zhou, 2014;Seto and Yoshida, 2014). The KDACs are subdivided into four classes (I, II, III/sirtuins, IV). Class III KDACs, also known as sirtuins, are homologous to yeast Sir2 and require nicotinamide adenine dinucleotide (NAD^+^) as cofactor for their catalytic activity (Seto and Yoshida, 2014). Sirtuins are conserved from bacteria to human (Brachmann et al., 1995). These are multifunctional enzymes involved in several biological processes, as gene expression regulation, DNA repair, metabolism and oxidative stress response (Choi and Mostoslavsky, 2014). The number of genes coding for sirtuins varies depending on the species; while human has 7 sirtuins, bacteria has only one. Three sirtuins are present in *T. brucei* (TbSir2rp1-3) and *Leishmania* spp. (LmSir2rp1-3), while only two are described in *T. cruzi* (TcSir2rp1 e 2) (Alsford et al., 2007;Tavares et al., 2008;Moretti et al., 2015;Ritagliati et al., 2015;Vergnes et al., 2016).

The TcSir2rp1 and TcSir2rp3 sirtuins have cytosolic and mitochondrial localization, respectively (Moretti et al., 2015;Ritagliati et al., 2015). Overexpression of TcSir2rp3 improves epimastigote multiplication and epimastigote differentiation to infective metacyclic forms. Moreover, increases amastigote intracellular multiplication, phenotype that is negatively affect when an inactive version of TcSir2rp3 was overexpressed or specific sirtuin inhibitors, such as salermide, are used (Moretti et al., 2015).

In this work, we investigated the involvement of *T. cruzi* mitochondrial sirtuin in the regulation of oxidative stress response of the parasite. We investigated whether overexpression of TcSir2rp3 increases parasite resistance to oxidative stress and if the mitochondrial superoxide dismutase A could be regulated by TcSir2rp3. We further verified the enzymatic activity of recombinant forms of TcSODA to investigate the effects of acetylation and looked by molecular dynamic analysis the predicted effects of acetylation on the enzyme activity. Our results indicate for the first time a direct effect of lysine acetylation in the regulation of antioxidant enzymes in protozoan parasites.

## Materials and Methods

### Cloning of native and mutated versions of the TcSOD-A protein

To obtain the native version of *T. cruzi* SOD-A protein (TcCLB.509775.40) (http://tritrypdb.org), the gene was PCR amplified from genomic DNA of *T. cruzi* Y strain using the oligonucleotides TcSODA-Fow (5’-TCTAGA<u>CATATG</u>TTGAGACGTGCGGTGAA-3’), TcSODA-Rev (5’-CC<u>GGATCC</u>TTATTTTATGCCTGCGCATGCAT-3’). The underlined residues represent cloning sites, for NdeI and BamHI enzymes, respectively. To generate mutated TcSODA proteins, we used two rounds of PCRs. In the first round, for TcSODA-K44Q and TcSODA-K97Q, where the lysine (K) residues were replaced by glutamine (G), by changes in the nucleotides A130 and A289 to cytosines, respectively. For TcSODA-K97R, the K was substituted by arginine (R) replacing the nucleotide A289 by guanine. In the second round of PCR, we used the same primers described above to obtain the native version of TcSOD-A. The oligonucleotides used to obtain the mutated TcSOD-A versions are listed in Table S1. The native and mutated TcSOD-A fragments were digested with NdeI and BamHI and cloned into the expression vector pET28a (New England Biolabs), using the same restriction sites. Clones were selected and submitted to sequencing reactions to confirm the desired mutations.

### TcSOD-A protein heterologous expression and purification

The pET28a vectors bearing native or mutated versions of TcSODA (K44Q, K97R and K97Q), were transformed into BL21 bacteria (DE3) (New England Biolabs); clones were selected and used for protein heterologous expression. For expression bacteria were growth at 37°C in LB medium containing 50 μg/mL kanamycin and induced by addition of 0.1 mM IPTG at 0.6-0.8 OD at absorbance of 600 nm and incubated for an additional 4 h. The cells were collected by centrifugation at 5,000 × g for 10 minutes, washed once with 20 mM Tris-HCl, pH 8.0; and the pellets were resuspended in lysis buffer (200 mM NaCl, 5% Glycerol, 5 mM 2-mercaptoethanol, 25 mM Hepes-NaOH, pH 7.5 and proteases inhibitors), and lysed using the French Press apparatus to obtain the soluble fractions of proteins. The soluble fractions were incubated with Ni-NTA Agarose resin (QIAGEN), previously equilibrated with equilibrium buffer (200 mM NaCl, 5% glycerol, 5 mM 2-β-mercaptoethanol, 25 mM imidazole, 25 mM HEPES-NaOH, pH 7.5). After 30 minutes under agitation at 4 °C the resin was washed 4 times with 5 ml of wash buffer (200 mM NaCl, 5% glycerol, 5 mM 2-β-mercaptoethanol, 50 mM imidazole, 25 mM HEPES-NaOH pH 7.5) and proteins were eluted with 4 mL of elution buffer (200 mM NaCl, 5% glycerol, 5 mM 2-β-mercaptoethanol, 250 mM imidazole and 25 mM HEPES-NaOH, pH 7.5). The quality of the purification was confirmed by SDS–PAGE gel electrophoresis and staining with Coomassie Blue for further use.

### TcSOD-A enzyme activity assays

The activity of superoxide dismutase was determined using the oxidation of 4-Nitro blue tetrazolium chloride (NBT) (Sigma-Aldrich) method, as described in (Ewing and Janero, 1995). To measure the activity of purified recombinant TcSODA proteins (native and mutated versions), proteins were initially dialyzed against the superoxide dismutase reaction buffer (50 mM KPO_4_, 0.1 M EDTA, pH 7.4), concentrated using Amicon Ultra – 10 K columns (Merck Millipore) and quantified with Pierce^™^ BCA Protein Assay Kit (Thermo Fischer Scientific). A total of 10 μg of dialyzed proteins (native and mutant versions) were incubated with 1x reaction buffer (50 mM KPO_4_, 0.1 M EDTA, pH 7.4), 50 μM NBT (Sigma-Aldrich), 78 μM NADH (Sigma-Aldrich) and 3.3 μM of phenazine (Sigma-Aldrich), in a 96-well plate. The enzymatic activity was determined by the detection of NBT oxidation at absorbance of 560 nm using the SpectraMax M3 equipment (Molecular Devices), every 30 seconds during 5 minutes. When indicated 5 μg of purified TcSODA was incubated in 50 mM Tris-HCl pH 7.4 with different concentrations of anhydride acetic at 1 μM; 2.5 μM; 5 μM; 10 μM) at room temperature for 30 minutes and the samples were dialyzed against 50 mM Tris-HCl pH 7.4.

The superoxide dismutase activity in *T. cruzi* extracts of wild-type, Sir2rp3-WT and Sir2rp3-MUT parasites, previously described in (Moretti et al., 2015), was performed using epimastigote forms at exponentially growing phase. The parasites were collected by centrifugation, washed in PBS 1X and total protein extracts obtained by freezing and thaw in PBS 1X and 1x Complete-EDTA free protease inhibitors and submitted to dialysis against SOD reaction buffer (50 mM KPO_4_, 0.1 M EDTA, pH 7.4), and further used to activity assays as described above.

### Parasite cultivation and growth curve experiments

For this work we used *T. cruzi* Y strain wild-type and two cell lines overexpressing the mitochondrial sirtuin, TcSir2rp3, in its native (Sir2rp3-WT) and inactive (Sir2rp3-MUT) form, generated previously (Moretti et al., 2015). Epimastigote forms of all cell lines were cultivated in liver infusion tryptose (LIT) medium supplemented with 10% fetal bovine serum (FBS) at 28 °C, and for transfectant parasites 500 μg/mL of geneticin (G418) was added to the medium.

The effect of hydrogen peroxide (H_2_O_2_) was determined in epimastigotes inoculated at 2×10^7^ parasites/mL in LIT medium containing 100 μM of H_2_O_2_ at 28°C. After 20 minutes the parasite cells were collected by centrifugation and resuspended in fresh LIT medium without H_2_O_2_, and parasite numbers was measured after 24 h with Neubauer chamber. In the experiments with menadione, 5×10^6^ parasites/mL epimastigote forms of all cell lines were inoculated in LIT medium in the presence of 10 μM of the drug and the parasite growth was determined after 48 h. The experiments using nicotinamide were carried out as described above, but with the addition of 10 mM of nicotinamide in the described conditions.

To determine the resistance of cell lines against benznidazole (BZD) and nifurtimox (NFX), 2×10^6^ parasites/mL epimastigote forms were inoculated in LIT medium in 24-well plates, in the presence of increasing drug concentrations and kept growing at 28 °C for four days. After the incubation time, parasite cell growth was determined by counting the cells with Neubauer chamber, and BZD/NFX resistance was calculated by comparing parasite growth in the absence versus parasite growth in the presence of drugs. The experiments in the presence of salermide were performed as described above, with addition of 25 μM of the inhibitor.

### Quantification of reactive oxygen species (ROS) levels

To determine the levels of reactive oxygen species (ROS), the parasites were initially exposed to 25 μg/mL of BZD or 15 μM of NFX for 24h in LIT medium at 28°C. These concentrations correspond to twice the EC_50_ of these drugs in the employed *T. cruzi* lineage previously determined. After incubation, 1.5×10^7^ parasites were collected from each cell line, and incubated for 30 minutes with 5 μM of CellROX Deep Red (Thermo Fisher Scientific #C10422) or CellROX Green (Thermo Fisher Scientific #C10444) reagents, to measure the cytoplasmic and mitochondrial ROS, respectively. Then, the parasites were washed twice with PBS 1X, and the fluorescence measured using flow cytometry with BD Accuri^™^ C6 Flow Cytometer (BD Biosciences) equipment. As a positive control of the experiment, parasites were exposed to 100 μM H_2_O_2_ for 20 minutes and washed once with PBS 1X before incubation with both reagents. All experiments were carried out in triplicate.

### Protein immunoprecipitation assays

Approximately 1×10^9^ cells were collected, washed once in PBS, and used to obtain total protein extracts of exponential cultures (1-1.5 × 10^7^ cells/mL) of the epimastigotes grown in LIT medium at 28 °C. The parasites were lysed in lysis buffer (20 mM Tris-HCl, pH 8.0; 137 mM NaCl; 2 mM EDTA; 1% Triton X-100; 25 mM nicotinamide; 25 mM sodium butyrate and protease inhibitors), and the soluble fraction was collected after centrifugation for 10 minutes, at 13,000 *g* at 4 °C. The soluble fraction was incubated with Dynabeads Protein A (Thermo Fisher Scientific), previously coupled with anti-TcSOD-A or anti-TcSir2rp3 antibodies, obtained during this work in our laboratory. After 2 h of incubation, samples were washed three times with washing solution (10 mM Tris-HCl pH 7.4; 1 mM EDTA; 1 mM EGTA; 150 mM NaCl; 1% Triton X-100 and 1x Complete-EDTA free protease inhibitors), followed by elution of the immunoprecipitated proteins with 0.2 M of glycine pH 2.0. The samples were then neutralized and used in the western blot experiments to detect the protein interactions with specific antibodies.

### Western Blot experiments

Protein extracts were obtained by lysing the parasites with lyses buffer (PBS 1X, 1% of Triton X-100, 0.5 mM of phenyl-methylsulfonyl fluoride (PMSF) and 1x Complete-EDTA free protease inhibitors. The samples were mixed with 2x Laemmli Buffer (Sigma-Aldrich) and used in SDS-PAGE electrophoresis with 12.5% polyacrylamide gels. After electrophoresis, the proteins were transferred to nitrocellulose membranes (BioRad Laboratories) using the Trans-blot SD Semi-Dry system (BioRad Laboratories). The membranes were incubated with blocking solution (1x PBS, 0.01% Tween 20 and 5% skimmed milk powder) for 1h under agitation, followed by incubation with specific primary antibodies diluted in blocking solution; anti-TcSir2rp3 (1:2,500); anti-TcSODA (1:5,000); anti-Histone H3 (1:10,000); anti-aldolase (1:5,000), anti-acetyl-lysine (1:2,000) for 1h under agitation at room temperature. The membranes were washed three times with PBS 1X containing 0.01% Tween 20, and incubated with secondary antibodies IRDye^®^ 800CW Goat anti-Rabbit IgG Secondary Antibody or IRDye^®^ 700CW Goat anti-Mouse IgG Secondary Antibody diluted 1:10,000 in blocking buffer (Li-Cor), during 1h at room temperature. The membranes were washed three times with washing solution and revealed using Odyssey Li-Cor equipment. The same procedure was used for western blot analysis with samples obtained in protein immunoprecipitation assays or for samples from acetic anhydrite assays.

The primary antibody anti-TcSODA and anti-TcSir2rp3 were house made during this work; anti-aldolase was obtained previously in our laboratory (Barbosa Leite et al., 2020), anti-histone H3 was kindly provided by Prof. Dr. Christian Janzen from University of Würzburg, Germany and anti-acetyl-lysine was purchased from Millipore (#ab80178).

### Molecular Dynamics Simulations of TcSODA

The crystallographic structure of TcSODA was retrieved from the Protein Data Bank (PDB id: 4H3E) (Phan et al., 2015), preserving only the coordinates of Fe, and submitted to Molecular Dynamics (MD) simulations using GROMACS v.2020.6 under the CHARMM36m force field (Huang et al., 2017). The protonation state of TcSODA was determined according to the PROPKA3 webserver (Olsson et al., 2011). To study the effect of the acetylation in K97, TcSODA was used as a monomer in both native (WT) and acetylated (K97Ace) states. Initially, each configuration of TcSODA was centered in a rhombic dodecahedric box of 12 Å distant from the farthest atom in XYZ directions. Further, each system was solvated and equilibrated with 0.1 M of NaCl, with some additional ions added to reach a system with zero charge. As previously described, iron superoxide dismutases show a Fe^3+^ bound to the protein at the first step of the enzymatic process, thus, the Fe3^+^ topology was obtained from a previous work in order to reproduce the same conformational state of the enzyme (Li et al., 2015). Each system was minimized using the Steepest Descent algorithm until reach an energy gradient below 100 kJ/mol/nm^2^. Following, each system was submitted to an NVT ensemble generating the initial velocities randomly following a Maxwell-Boltzman distribution at 300 K during 1 ns, monitoring the temperature using the V-Rescale thermostat (Bussi et al., 2007) with a time constant of 0.1 ps. Then, an NPT ensemble of 1 ns was applied maintaining the pressure of the system at 1 bar using the Berendsen barostat (Eslami et al., 2010). Both steps were performed restraining the backbone atoms of the protein and Fe^3+^ with a force constant of 1000 kJ/mol/nm^2^. Finally, an unconstrained MD of 200 ns was applied using the Nose-Hoover thermostat (Hoover, 1985) and Parrinello-Rahman barostat. Non-bonded interactions were calculated considering atoms within 10 Å using the PME method, with a switching-force function between 10 and 12 Å. Three independent replicas for both wildtype and acetylated systems were performed collecting frames every 10 ps.

Two structural analyses were performed to study the acetylation effect on TcSODA: (i) the volume occupancy of the sidechain nitrogen atom of K97 or K97Ace was calculated using the VolMap tool at a resolution of 1 Å and Isovalue of 0.25; and (ii) the minimal distance of the sidechain nitrogen atom of K97 to the sidechain oxygen atoms of D94 or E96 residues of TcSODA. Both analyses were performed using VMD (Humphrey et al., 1996).

### Ethics Committee

This work was carried out in accordance with the rules of the Research Ethics Committee (CEP) of the Universidade Federal de São Paulo - UNIFESP. The experimental protocol was approved by the CEP under number 6383311017.

## Results

### Mitochondrial *T. cruzi* sirtuin overexpression increases parasite resistance to H_2_O_2_ and menadione

As sirtuins regulate the levels of protein acetylation, and in *T. cruzi*, several proteins involved in redox reactions are acetylated (Moretti et al., 2015), we initially explored how TcSir2rp3 could impact parasite responses to oxidative stress. For this, we employed the *T. cruzi* cell lines overexpressing the mitochondrial sirtuin, TcSir2rp3, in its native (Sir2rp3-WT) and inactive version (Sir2rp3-MUT). These two cell lines expressed similar levels of TcSir2rp3 which were about three times than the endogenous protein in the non-transfected line, as demonstrated in (Bussi et al., 2007).

The different epimastigote cell lines were subjected to oxidative stress by hydrogen peroxide (H_2_O_2_) and menadione and the recovering after stress was determined. In addition to growth faster (Moretti et al., 2015) the Sir2rp3-WT cell line recovered faster from the oxidative stress caused by H_2_O_2_ compared to WT or Sir2rp3-MUT parasites (Figure 1A). In the case of menadione, a better recovery was detected for Sir2rp3-WT but a large reduction in parasite growth was observed in Sir2rp3-MUT parasites compared to WT cell line (Figure 1C).

**Figure 1.**
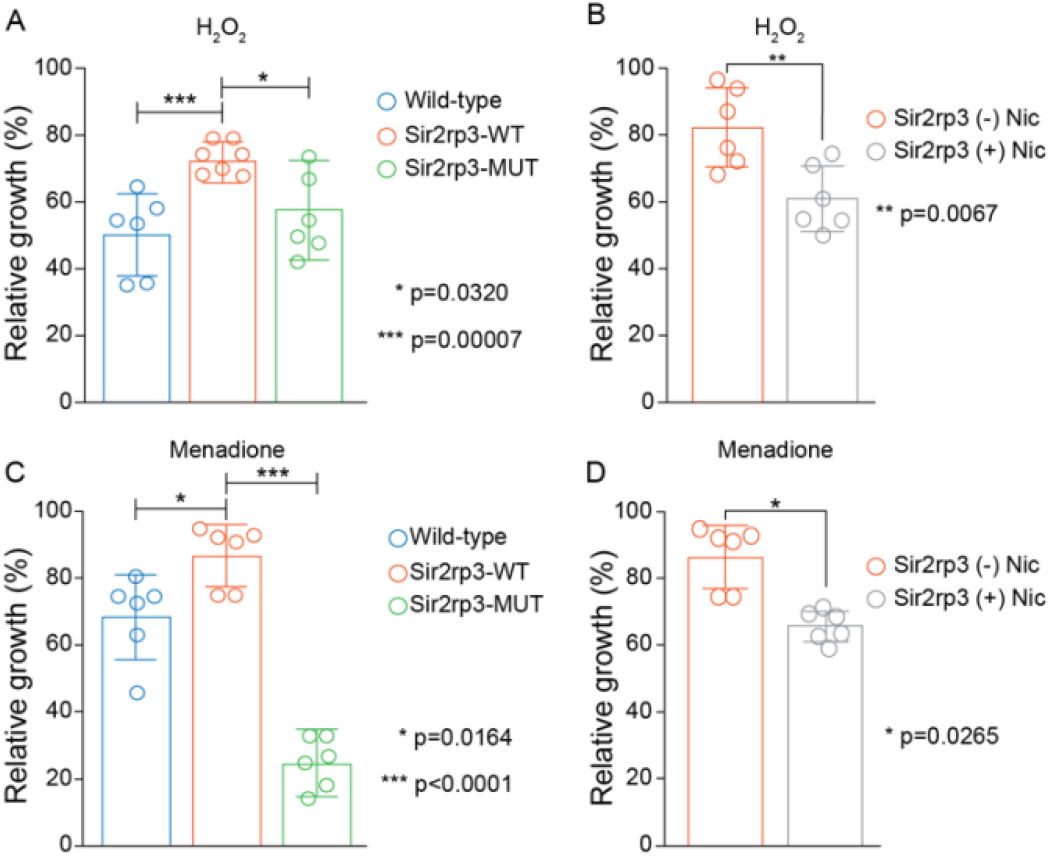
Overexpression of mitochondrial TcSir2rp3 improves parasite recovery after oxidative stress. **(A and B)**. Epimastigotes forms of WT (blue bars), Sir2rp3-WT (orange bars) and Sir2rp3-MUT (green bars) were submitted to oxidative stress with 100 μM H_2_O_2_ and 10 μM menadione and recovering was quantified. **(C and D)**. Sir2rp3-WT (orange bars) recover was analyzed in the assays performed in the presence of nicotinamide (Nic) (grey bars) compared to assays performed without the inhibitor (orange bars). All experiments were performed in triplicate. Statistical analyzes were performed using the student test *t*. *(p=0.0320); ***(p=0.00007) in A; **(p=0.0067) in B; *(p=0.00164) and ***(p<0.0001) in C; *(p=0.0265) in D.

To investigate if the phenotype observed for Sir2rp3-WT overexpressor was directly related to enzyme activity we performed the same assays in the presence of nicotinamide, a sirtuin inhibitor. A reduction in the recovering phenotype was detected in the presence of nicotinamide in the Sir2rp3-WT cells, compared to those cultivated in the absence of nicotinamide (Figure 1B and D).

### TcSir2rp3 overexpression increases parasite resistance to BZD and NFX

Based on the findings that TcSir2rp3 improves *T. cruzi* recovering after H_2_O_2_ and menadione stress and in the fact that the two drugs available for treatment of Chagas disease, nifurtimox (NFX) and benznidazole (BZD), acts through oxidative damaged on the parasites, we decided to evaluate the effect of TcSir2rp3 overexpression in BZD and NFX resistance.

For that, epimastigotes of WT, Sir2rp3-WT and Sir2rp3-MUT were cultivated in the presence of increasing concentrations of BZD and NFX, and their survival was evaluated after four days of growing compared to mock-treated parasites. The relative growth was higher for the Sir2rp3-WT parasites showing that Sir2rp3 induces resistance to both drugs, NFX and BZD, compared to WT and Sir2rp3-MUT cells (Figure 2A and B). More important, the resistance phenotype observed for Sir2rp3-WT cells was reverted when the assays were performed in the presence of salermide, another specific sirtuin inhibitor (Figure 2). We also noticed difference responses for the Sir2rp3-MUT with respect of resistance to BZD and NFX.

**Figure 2.**
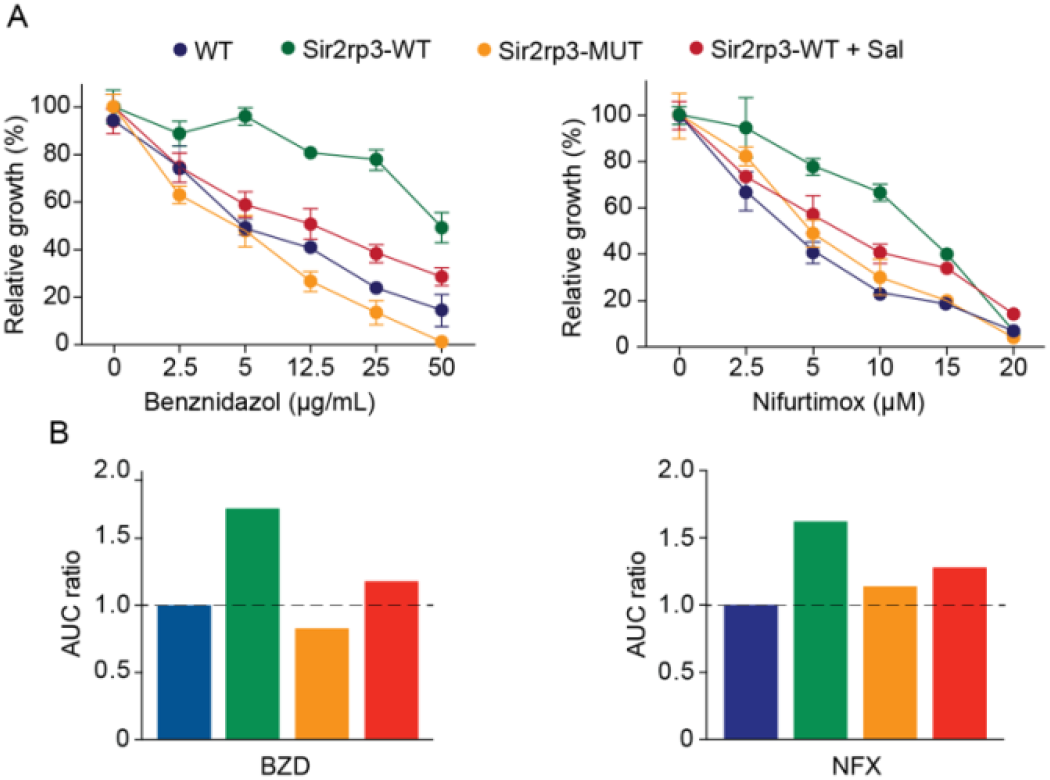
TcSir2rp3 increases parasite resistance to BZD and NFX. Epimastigotes of the indicated lines were incubated with BZD **(A and B)** or NFX **(C and D)** and the relative growth determined by the percentage of parasites in the presence of absence of each compound. All the values are mean ± SEM of triplicates. To better visualize the effects panels **(B and D)** show the area under the curve (AUC) of the data shown in A and C.

### Cytosolic and mitochondrial ROS levels are lower in Sir2rp3-WT parasites after BZD and NFX treatment

To know if the BZD and NFX resistance observed in the Sir2rp3-WT parasites was related to reduction in the ROS levels, WT, Sir2rp3-WT and Sir2rp3-MUT epimastigotes were cultivated in the presence of 25 μg/mL of BZD or 15 μM of NFX, corresponding to twice the EC_50_ of these drugs in the employed *T. cruzi* strain, for 24 h and the levels of cytosolic and mitochondrial ROS were further quantified. Lower levels of ROS were detected in the Sir2rp3-WT cells compared to WT and Sir2rp3-MUT parasites (Figure 3), especially in the BZD treated parasites. In contrast, the ROS levels observed for Sir2rp3-MUT cells were higher than the WT parasites, suggesting a deleterious effect caused by the overexpression of an inactive TcSir2rp3 form.

**Figure 3.**
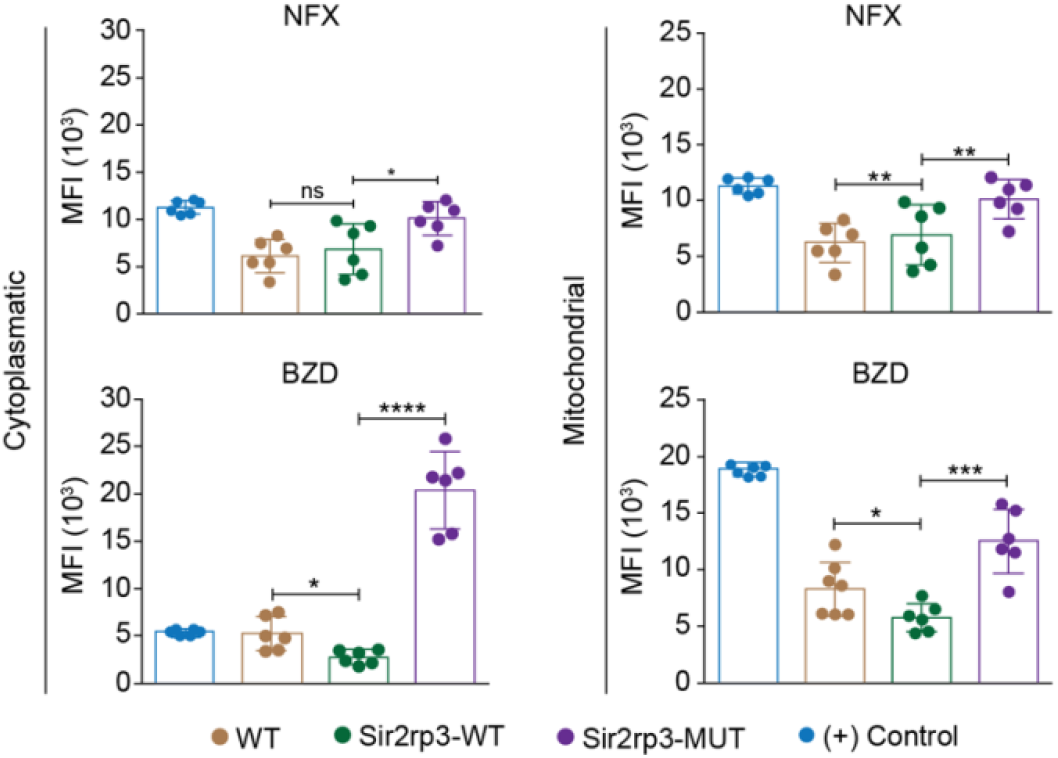
Mitochondrial sirtuin decreases ROS levels after BZD and NFX treatment. Sir2rp3-WT epimastigote presented lower cytosolic and mitochondrial ROS levels after BZD and NFX treatment compared to WT and Sir2rp3-MUT parasites. Significantly increase of ROS levels were observed in Sir2rp3-MUT. WT parasites submitted to H_2_O_2_ were used as positive control (+). All experiments were performed in triplicate. Statistical analyzes were done using the student test *t*. BZD analysis *(p=0.0105); ***(p<0.0003); ****(p<0.0001). NFX analyzes *(p=0.0067); **(p=0.0019).

### TcSir2rp3 overexpression does not affect TcSODA protein expression

To investigate the mechanisms involved in the resistance to H_2_O_2_, menadione, BZD and NFX, observed in the parasites overexpressing TcSir2rp3, we decided to compare the protein expression level of a key antioxidant enzyme, mitochondrial superoxide dismutase (TcSODA), among epimastigotes from WT, Sir2rp3-WT and Sir2rp3-MUT. No large differences in protein expression levels were observed for TcSODA among the cell lines (Figure 4A and B).

**Figure 4.**
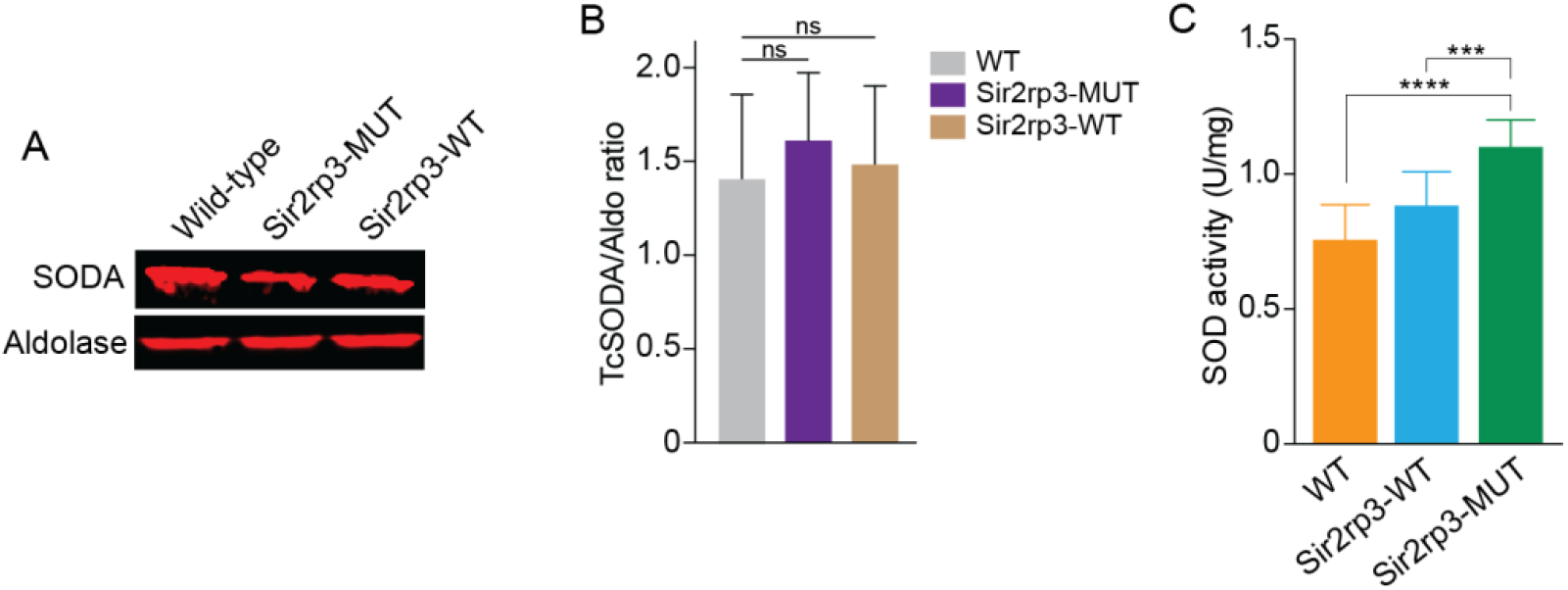
TcSir2rp3 overexpression increase superoxide dismutase activity in *T. cruzi*. **(A) A and B**. Comparative analyses of protein expression of antioxidant enzymes in the wild-type, Sir2rp3-WT and Sir2rp3-MUT parasites. No significative differences were observed among the cell lines for all enzymes analyzed. Aldolase was used as loading control. TcSODA levels were determined for all cell lines from a triplicate experiment. **(C)** SOD activity of total extracts of parasites. The assays were carried out in triplicate. Statistical analyzes were performed using the student test *t*. *** (p=0.0027); ****(p<0.0001).

### Superoxide dismutase activity increases after TcSir2rp3 overexpression

As human MnSOD activity was demonstrated to be negatively regulated by acetylation and that mitochondrial sirtuin SIRT3 is responsible to promote its deacetylation (Chen et al., 2011), we decided to measure superoxide dismutase activity in the parasites overexpressing TcSir2rp3.

Therefore, we measured the SOD activity in total protein extracts of WT, Sir2rp3-WT and Sir2rp3-MUT epimastigotes. We observed an increase of approximately 40% in superoxide dismutase activity in Sir2rp3-WT protein extracts compared to those from WT and Sir2rp3-MUT (Figure 4C). This increase was not due to augmented protein expression as western blot analyzes using anti-TcSOD-A antibody showed similar levels of TcSODA among the parasites used in the activity assays (Figure 4A and B). This result suggests that TcSODA activity might be higher in Sir2rp3-WT parasites, and this could be associated to decreased levels of TcSODA acetylation promoted by sirtuin activity.

### TcSir2rp3 and TcSOD-A interacts in vivo

The set of experiments performed before suggests that TcSir2rp3 might be involved in oxidative stress response in *T. cruzi* by regulating the acetylation level of TcSODA. To gain insight in the in the regulatory function of TcSir2rp3 above TcSODA activity, we investigated the interaction of these two enzymes *in vivo* by protein immunoprecipitation assays.

Total protein extracts from WT parasites were obtained and immunoprecipitated with anti-TcSODA and samples submitted to western blot analyzes with anti-TcSODA and anti-TcSir2rp3. As shown in Figure 5, TcSir2rp3 and TcSOD-A were detected in the immunoprecipitated fraction, indicating that these proteins interact with each other in the parasite. The negative control of the experiment was done through immunoprecipitation with an unrelated antibody, anti-aldolase, and no interaction was observed (Figure 5).

**Figure 5.**
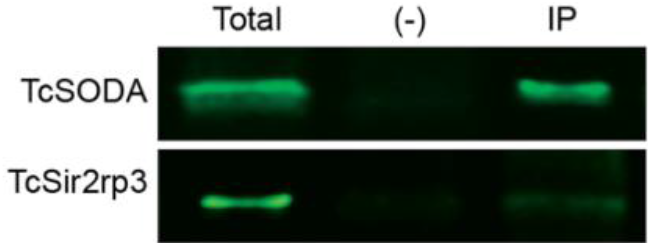
Interaction of TcSir2rp3 and TcSODA proteins. Immunoprecipitated samples with anti-TcSODA of total protein extracts of WT parasites were submitted to western blot analyzes with anti-TcSir2rp3 and anti-TcSODA. The two proteins, TcSir2rp3 and TcSODA, were detected in the immunoprecipitated fractions (IP). No signal was detected in the negative control samples (−) immunoprecipitated with anti-aldolase. Experiments were performed in triplicated and the figure is a representative of the results obtained.

### K97 acetylation negative regulates TcSOD-A *in vitro* activity

To further understand the function of lysine acetylation in the regulation of TcSOD-A activity, we submitted heterologous TcSOD-A in its native state (WT) to acetic anhydrite (AA) treatment, a chemical compound that acetylates any exposed lysine residues followed by TcSODA *in vitro* activity assays. Crescent concentrations of AA followed by western blot with anti-acetyl lysine antibody showed an increased in the acetylation levels of TcSODA heterologous protein (Figure S1A). In parallel AA treatment samples were used in the *in vitro* activity assays and we observed that acetylation by AA reduced TcSODA activity compared to non-treated samples (Figure S1B). This result suggested that acetylation might negatively impact TcSODA enzyme activity.

In the *T. cruzi* acetylome, two lysine-acetylated sites were identified in TcSODA, K44 and K97 (Moretti et al., 2018). To evaluate the effect of acetylation at K44 and K97 residues in the regulation of enzyme activity, we used purified heterologous protein of WT and mutated versions mimicking acetylation at K44 and K97 (K44Q and K97Q) or non-acetylated state at K97 (K97R), to measure the superoxide activity *in vitro* by the NBT oxidation method. The acetylation of K97 reduced TcSODA activity in about 25% compared to WT protein (Figure 8), which was not observed in the K97R that mimics non-acetylated state or K44Q that is located in a region distant from the tunnel of the catalytic site.

### MD simulations suggests that the K97 acetylation affects the salt-bridge pattern of TcSODA

To better understand how the acetylation pattern of TcSODA affects its ability to process superoxide, we performed MD simulations of the native (WT) and K97 acetylated (K97Ace) versions of TcSODA. The lysine 97 is placed at around 21 Å away from the catalytic site (Figure 7, left panel), and despite this acetylation reduces the enzymatic activity of TcSODA by ~25%. Initially, we analyzed the conformational changes locally around the residue in both WT and mutant, and detected that the sidechain nitrogen of TcSODA showed an evident difference in spatial occupancy comparing WT and K97Ace. The WT maintained the lysine crystallographic orientation, while the acetylated one presented its sidechain with a tendency to move away from that position (Figure 7, right panel). This behavior can be explained by the interaction with neighbor residues, in which K97 is able to interact to with two near acidic residues, named D94 and E96, while K97Ace tends to be repealed due to charge modification caused by the acetyl group.

**Figure 6.**
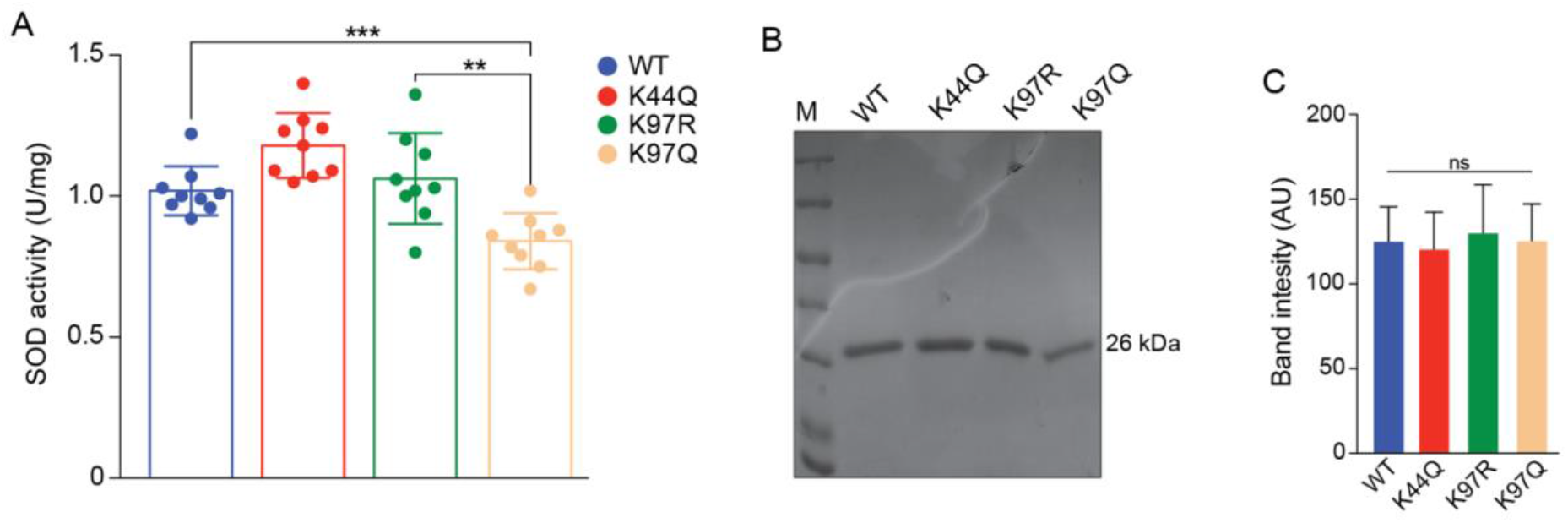
Superoxide dismutase activity of native and mutated versions of TcSOD-A. Purified heterologous proteins of TcSODA (native and mutated versions) were submitted to superoxide dismutase assays using NBT oxidation method. A reduction of about 25% was observed for K97Q protein, which mimics acetylation, compared to the native (WT) or K97R versions. No significant difference was detected for K44Q compared to WT or K97R. All analyzes were performed in triplicate. Statistical analyzes were performed using the student test *t*. **(p = 0.0028); ***(p = 0.0009). **B**. Comassie gel of TcSODA recombinant proteins used in the in vitro activity assays. **C**. Quantification of triplicate comassie gel of recombinant proteins used in the TcSODA activity assays. No significant differences were observed among the samples.

**Figure 7.**
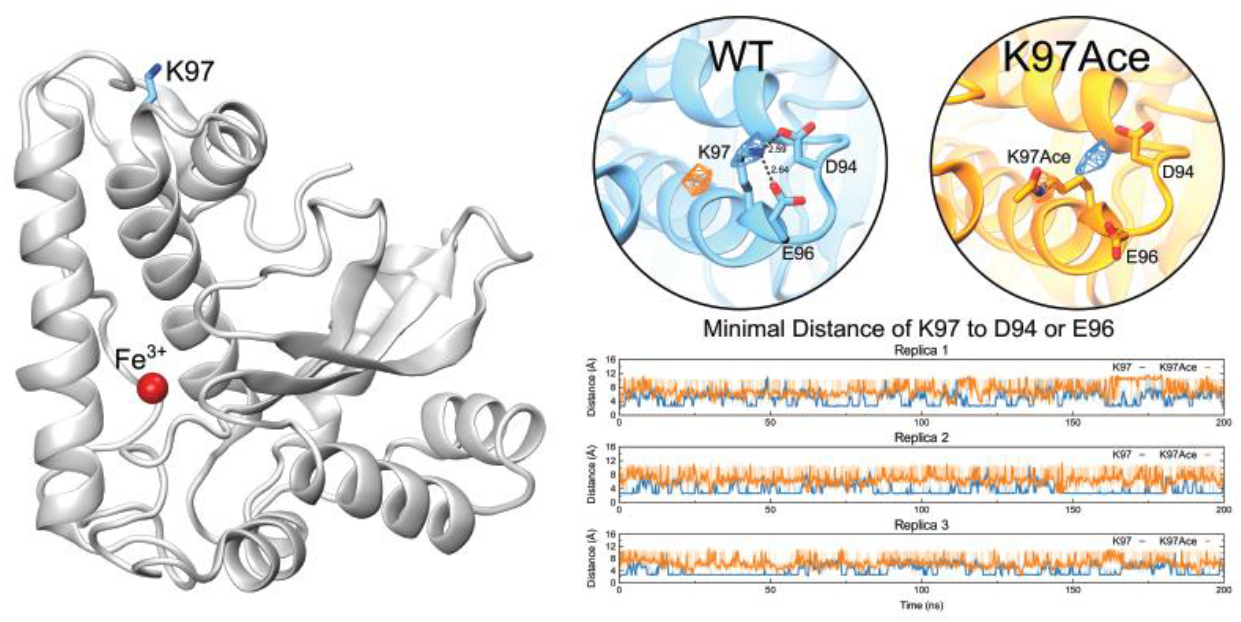
MD simulations reveals that K97 acetylation affects the salt-bridge pattern of TcSODA. The left panel shows TcSODA structure as cartoon, highlighting the position of K97 residue and Fe^3+^, shown as cyan sticks and as a red sphere, respectively. The right panel shows the orientation of K97 in respect to D94 and E96 in native (WT, in cyan) and acetylated (K97Ace, in orange) TcSODA, measured by the occupancy of the sidechain nitrogen atom of K97, showing that K97 is kept near these acidic residues whereas acetylation repeals it. This spatial change of K97 alters the salt-bridge formation between such residues, as shown by minimal distance measurements in all replicas.

To compare the dynamical behavior of K97 in WT or acetylated in respect of neighbor acidic residues, the minimal distance between the sidechain nitrogen of K97 to the sidechain oxygen atoms of D94 or E96 were calculated (Figure 7, right panel). It clearly showed how the WT lysine orients towards both residues, showing interactions at distances bellow 3.5 Å, which correspond to salt-bridge interactions, in 52.21% of the time, while the acetylated enzyme showed such interaction in only 1.46%. This shows that the acetylation caused a ~36-fold reduction of salt-bridges between K97 and D94 or E96 in TcSODA.

## Discussion

*T. cruzi* is exposed to different redox environments during its life cycle and adaptation to these conditions determines the success of the infection (Perez-Molina and Molina, 2018;Lidani et al., 2019). To deal with this situation the parasite has developed a sophisticated arsenal of antioxidant enzymes that help maintain redox homeostasis (Beltrame-Botelho et al., 2016). In this work, we showed for the first time that protein acetylation can regulate the oxidative stress response through the action of the sirtuin TcSir2rp3, a lysine deacetylase enzyme located in the mitochondria (Moretti et al., 2015). Here, we demonstrated that sirtuin deacetylation activity increased the resistance to different oxidants (H_2_O_2_, menadione, BZD and NFX) and reduced the levels of cytoplasmic and mitochondrial ROS agree with was observed in different organisms, suggesting an evolutionary conservation of the mechanisms involved in the regulation of oxidative stress response. We showed the mitochondrial SOD of the parasite is one of the targets of the sirtuin, as the specific activity of the enzyme is increased upon acetylation both in vitro and in vivo. Based on molecular modeling we suggest that acetylation of specific lysine residues negatively regulates TcSODA activity by affecting protein conformation, which causes changes in the active site.

Sirtuins are present from bacteria to humans and act regulating diverse cellular processes from different cellular compartments (Choi and Mostoslavsky, 2014). SIRT3 is the main mitochondrial sirtuin in humans playing essential role on the homeostasis of this organelle through the deacetylation of metabolic proteins, including those involved with ROS detoxification processes (Cooper and Spelbrink, 2008;Qiu et al., 2010). Also, it is known that SIRT3 knockout cells show high levels of ROS (Ahn et al., 2008;Tao et al., 2010). Our results demonstrating that parasites overexpressing the mitochondrial sirtuin, TcSir2rp3, increased parasite resistance to different oxidants compounds (H_2_O_2_, menadione, NFX and BZD) and reduced the levels of cytoplasmic and mitochondrial ROS are in agreement with was observed in mammals, suggesting an evolutionary conservation of the mechanisms involved in the regulation of oxidative stress response.

Although our data only provides evidence that TcSODA is regulated by acetylation, we cannot exclude that the resistance to oxidants, depends on the levels of other antioxidant enzymes by different mechanisms. Nevertheless, five *T. cruzi* antioxidant enzymes were identified acetylated at different sites (Figure 8 and Table S2), suggesting that acetylation might affect their activities as observed for antioxidant enzymes from other organisms (Narita et al., 2019). The regulatory function of lysine acetylation has been demonstrated for different antioxidant enzymes in humans, such as peroxiredoxins (PrxI and PrxII), superoxide dismutase 1 (SOD1) and mitochondrial MnSOD (Parmigiani et al., 2008;Chen et al., 2011;Lin et al., 2015).

**Figure 8.**
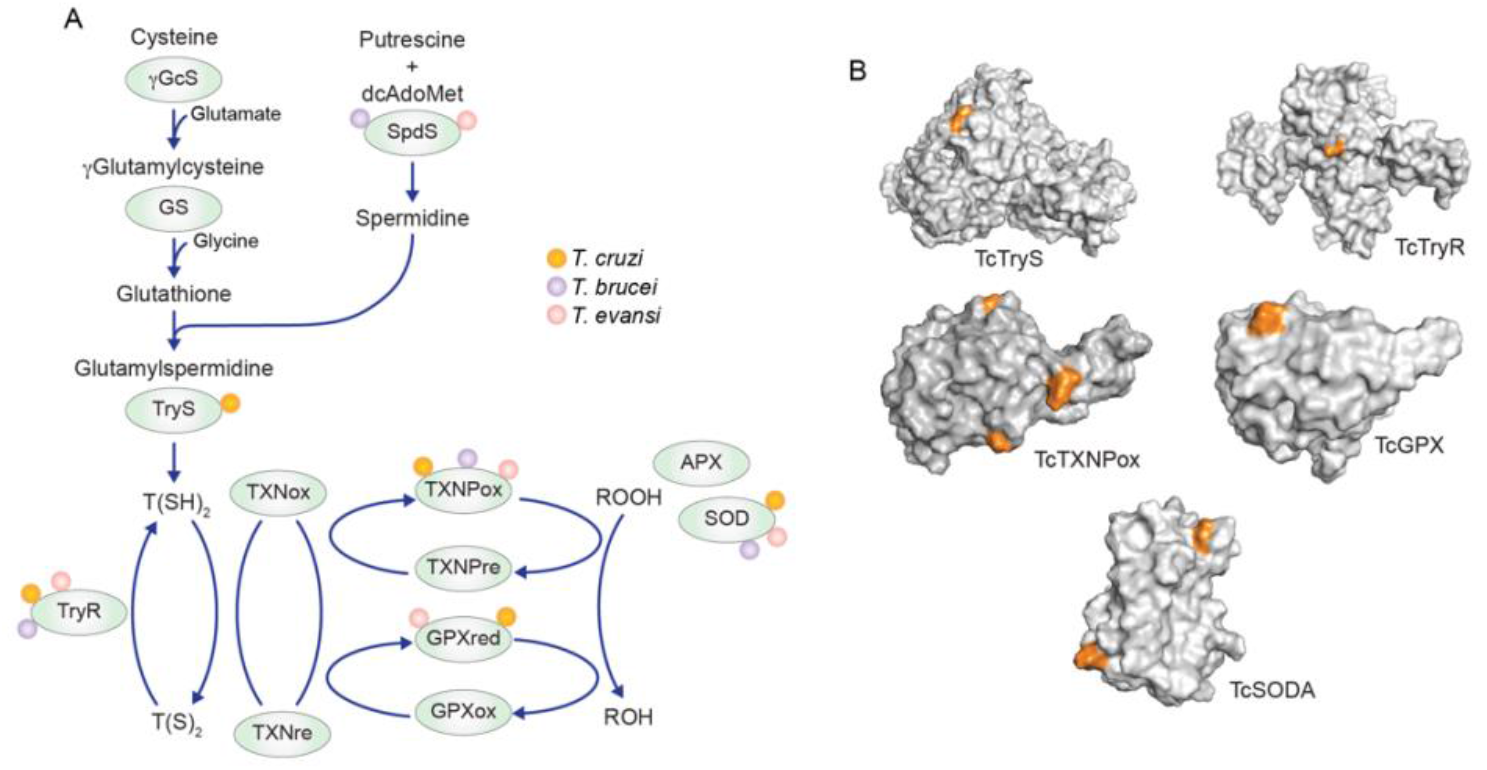
Antioxidant enzymes detected acetylated in *T. cruzi, T. brucei* and *T. evansi* acetylomes. **A**. Enzymes belonging to the tree main antioxidant pathways of trypanosomatids with one or more lysine sites detected acetylated. B. Distribution of lysine acetylated sites (orange) in the 3D protein structure of the five *T. cruzi* antioxidant enzymes detected in the acetylome of the parasite.

The SODs promote the dismutation of two molecules superoxide (O_2_-) generating H_2_O_2_ and O_2_, thus participating in the first line of defense of cells against oxidative stress (Wang et al., 2018). Our findings showing that overexpressing the native form of TcSir2rp3 increased superoxide activity is like what was found for human SIRT3, which reduced MnSOD acetylation increased the enzyme activity, contributing to the protection of cells against oxidative stress (Tao et al., 2010;Chen et al., 2011). Moreover, the direct interaction of TcSir2rp3 with TcSODA in the epimastigote forms, as observed in the protein immunoprecipitation assays, supports the notion that TcSODA activity could be directly modulated by TcSir2rp3. It will be interesting to demonstrate the TcSODA specific activity and determine the acetylation levels of the enzyme in the parasites overexpressing TcSir2rp3.

A comparison of the TcSODA and human MnSOD structures shows that the relative position of the acetylated K97 residue in *T. cruzi* is similar to that of the K68 residue in human MnSOD (Phan et al., 2015), suggesting that acetylation at K97 could regulates TcSODA activity. Through site-directed mutations we demonstrated that the replacement of K97 residue, the corresponding residue to K68 in humans, of TcSODA by glutamine, which mimics an acetylated lysine, causes decreased activity of the protein in vitro, as for human MnSOD (Chen et al., 2011;Lu et al., 2015). This effect was not seen when we replaced K97 with arginine, which mimics a deacetylated lysine, or when we perform a mutation in position K44, located in a region distant from K97. Thus, this data further supports the idea that TcSODA is regulated by acetylation. Moreover, TcSODA K97 residue is conserved in *T. brucei* and *T. evansi*, corresponding to K101, which was detected acetylated and is located at similar region in the protein 3D structure (Figure S1 and Table S2), indicating a conserved function in other trypanosomatids.

The direct effect of acetylation on the regulation of MnSOD activity has been proposed more than 30 years ago, and it was hypothesized that the tetrameric monomer complex of MnSOD would create a positively charged funnel structure that would facilitate the attraction of negatively charged O_2_- molecules to the catalytic site of MnSOD, and that the acetylation of lysines positioned at the funnel would cause electrostatic changes to the surface, thus affecting the activity of the enzyme (Benovic et al., 1983;Fridovich, 1983). Interestingly, we observed that K97 acetylation can induces particular conformational changes in TcSODA in respect of hydrogen bonding pattern to neighbor residues, specifically D94 and E96, suggesting a key participation of this residue to modulate the affinity to O_2_- by changing the charge availability on the surface of the enzyme. These findings show that lysine residues play a central role in TcSODA activity by sequestering negative charges of D94 and E96 in the native form, and exposing such negatively charged residues to the solvent when acetylated, modulating the affinity of the enzyme to O_2_-.

In addition, several studies demonstrates that replacement of MnSOD K68, which is located in a ring of 11 positively charged residues surrounding the active site (Borgstahl et al., 1992), by an residue mimicking an acetylated lysine or by an acetylated lysine, negatively affects the activity of the enzyme, and that this effect has a direct relationship with changes in the electrostatic potential of the protein, causing an increase in negative charges in the funnel region of the protein and decreasing the interaction of O_2_- with the catalytic site (Chen et al., 2011;Lu et al., 2015;Lu et al., 2017). Indeed, lysine acetylation modulates the O_2_- affinity by changing the global charge of the enzyme. In the present work we showed that lysine residues can be important to modulate the charge availability of TcSODA by sequestering negatively charged residues near the catalytic site by salt-bridge formation, suggesting that the molecular mechanism in free radical processing is likely more sophisticated than expected.

In conclusion, our results demonstrated for the first time the impact that lysine acetylation has in the regulation of redox state in trypanosomatids, contributing to better understand the mechanisms employed by *T. cruzi* to deal with the oxidant environmental during the progression of the infection. These observations open the opportunity to explore protein acetylation as potential drug target in this parasite, possible through combinatory therapy with the existent drugs. Moreover, this work helps to expand our knowledge about the function of lysine acetylation in the regulation of cellular processes in eukaryote organisms.

## Acknowledgements

We would like to thanks Claudio Rogério Oliveira for his technical help during the development of this work; Lucía Piacenza and Elton J. R. Vasconcelos for comments in the manuscript. This work was supported by the Fundação de Amparo à Pesquisa do Estado de São Paulo (FAPESP), grants 2018/09948-0 and 12/09403-8 to NSM; 15/22031-0 and 2020/07870-4 to SS; 14/03714-7 to VSN; by Conselho Nacional de Desenvolvimento Científico e Tecnológico (CNPq), grants 424729/2018-0 to NSM and 302883/2017-7 to MRMF; Instituto Nacional de Ciência e Tecnologia de Vacinas (INCTV) and scholarship 302972/2019-6 to S.S., and scholarship 165639/2017-2 to LSM; Coordenação de Aperfeiçoamento de Pessoal de Nível Superior scholarship 88887.506120/2020-00 to ACCNS.

## Supplementary files

**Sup. Figure 1.**
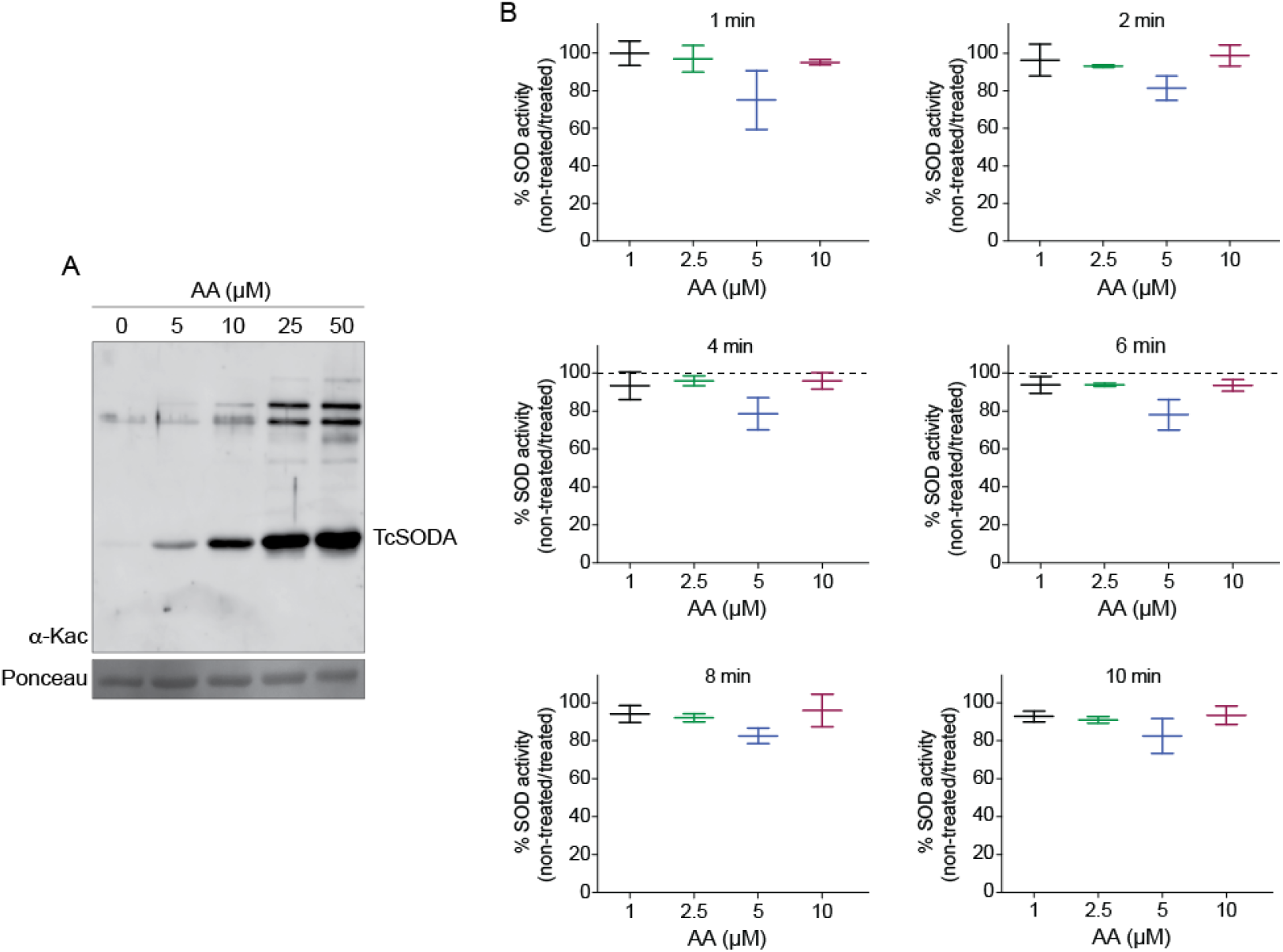
Effect of AA in the TcSODA in vitro enzyme activity. **A**. Treatment of TcSODA with AA increases protein acetylation. TcSODA-WT purified heterologous protein was submitted to treatment with different concentrations of AA and samples used in western blot analyses with anti-acetyl-lysine antibodies. The levels of lysine acetylation increased proportionally with AA concentrations. **B**. Treatment of TcSODA-WT with different concentrations of AA decreased the enzymatic activity *in vitro* compared to non-treated proteins. Different AA concentrations and time points were evaluated. All the experiments were performed in triplicate.

**Sup. Figure 2.**
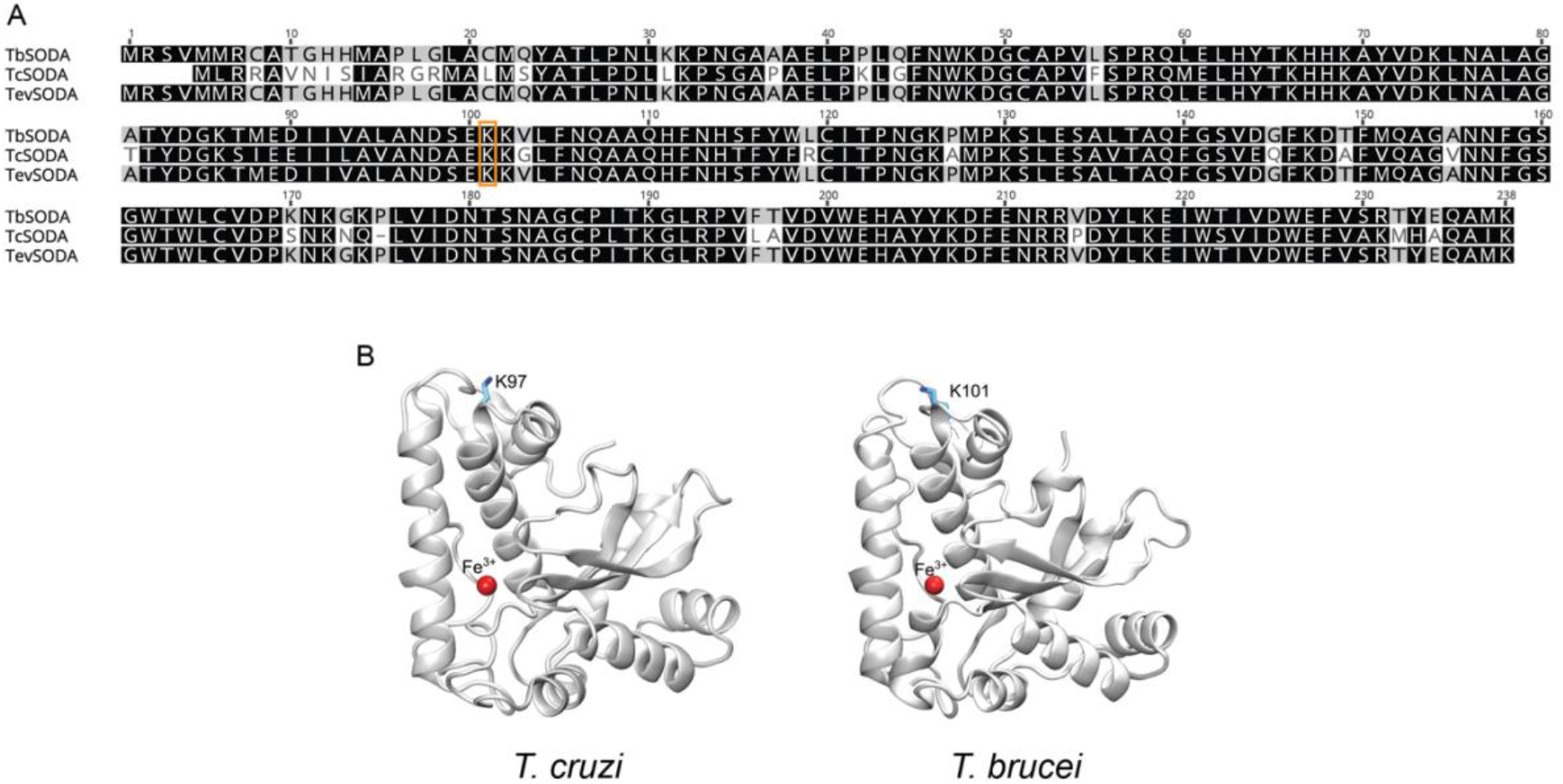
Conservation of K97 residue among *T. cruzi*, *T. brucei* and *T. evansi*. **A**. Amino acid sequence alignment of TcSODA (TcCLB.509775.40) and TbSODA (Tb927.5.3350) highlighting the conservation of lysine residue K97 between the two parasites. The conserved lysine residue is highlighted in orange and corresponds to K101 in *T. brucei* and *T. evansi*. **B**. Comparative 3D protein structure of TcSODA (PDB 4H3E) and TbSODA demonstrating that K101 is located at similar region of K97, suggesting a conserved regulatory mechanism in both parasites. *T. brucei* 3D protein structure was predicted using SwissModel https://swissmodel.expasy.org). *T. evansi* 3D structure was not included due to the high similarity with TbSODA protein.

**Table S1.**
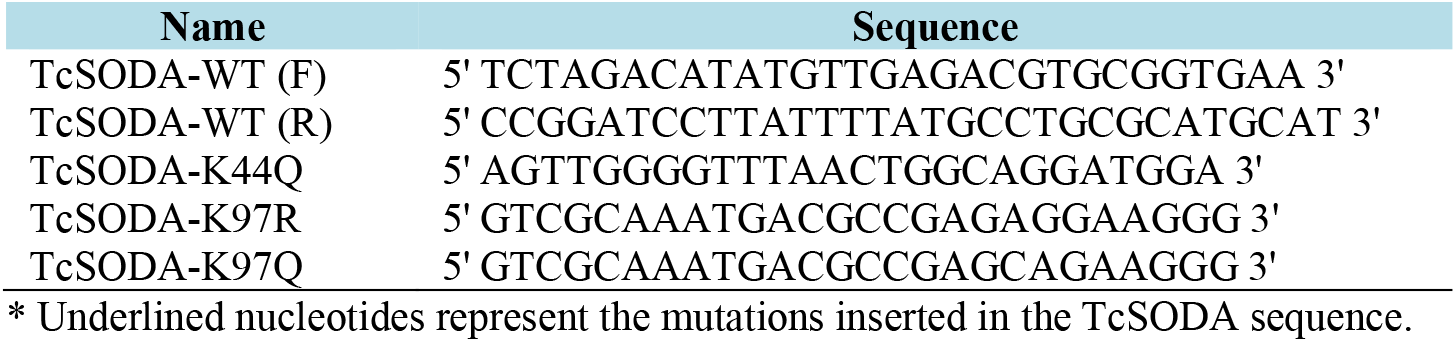
Oligonucleotides used to generate heterologous TcSODA

**Table S2.** Acetylated AD enzymes detected in *T. cruzi, T. brucei* and *T. evansi* acetvlomes

| *T. cruzi* |  |  |
| --- | --- | --- |
| TritrypDB ID | Gene name | Kac sites detected |
| TcCLB.509099.50 | Trypanothione synthetase | 604 |
| TcCLB.484299.10 | Trypanothione reductase | 147 |
| TcCLB.509775.40 | iron superoxide dismutase A, mitochondrial | 44; 97 |
| TcCLB.503899.130 | Glutathione peroxidase | 98 |
| TcCLB.487507.10 | Tryparedoxin peroxidase | 7; 64; 120; 168 |
| *T. brucei* |  |  |
| TritrypDB ID | Gene name | Kac sites detected |
| Tb927.11.15910 | iron superoxide dismutase | 22; 31; 138 |
| Tb927.11.15820 | iron superoxide dismutase C, mitochondrial | 75 |
| Tb927.11.15020 | iron superoxide dismutase | 39 |
| Tb927.5.3350 | iron superoxide dismutase A, mitochondrial | 66; 74; 101 |
| Tb927.7.1130 | trypanothione/tryparedoxin dependent peroxidase 2 | 43; 55; 89; 91; 115 |
| Tb927.8.1990 | peroxidoxin | 56; 64; 139; 165; 197; 219; 224 |
| Tb927.10.10390 | trypanothione reductase | 99; 144; 388 |
| Tb927.9.7770 | spermidine synthase | 220; 222; 273; 281 |
| Tb927.9.5860 | tryparedoxin peroxidase | 27; 28; 108; 120; 161; 168 |
| Tb927.7.4000 | glutathione synthetase | 223 |
| *T. evansi* |  |  |
| TritrypDB ID | Gene name | Kac sites detected |
| TEV003095.1 | iron superoxide dismutase, putative | 22; 31; 66; 74; 101; 135 |
| TEV006051.1 | glutathione peroxidase-like protein 3, putative | 43; 55; 57; 89; 91; 115 |
| TEV000603.1 | Chain A, Trypanothione Reductase | 60; 99; 144; 266 |
| TEV001721.1 | spermidine synthase, putative | 38; 222; 273; 281 |
| TEV004742.1 | tryparedoxin peroxidase | 64; 130; 139; 165; 197; 219; 224 |
| TEV006005.1 | tryparedoxin peroxidase | 27; 28; 120; 161; 168 |
| TEV006051.1 | glutathione peroxidase-like protein 3, putative | 43; 55; 57; 89; 91; 115 |

